# Comparative genomics of the world’s smallest mammals reveals links to echolocation, metabolism, and body size plasticity

**DOI:** 10.1101/2024.04.28.591546

**Authors:** Marie-Laurence Cossette, Donald T. Stewart, Aaron B.A. Shafer

## Abstract

Originating 30 million years ago, shrews (Soricidae) have diversified into around 400 species worldwide. Shrews display a wide array-array of adaptations, with some species having developed distinctive traits such as echolocation, underwater diving, and venomous saliva. Accordingly, these tiny insectivores are ideal to study the genomic mechanisms of evolution and adaptation. We conducted a comparative genomic analysis of four shrew species and 16 other mammals to identify genomic variations unique to shrews. Using two existing shrew genomes and *de novo* assemblies for the maritime (*Sorex maritimensis*) and smoky shrew (*S. fumeus*), we identified mutations in conserved regions of the genomes, also known as accelerated regions, gene families undergoing significant expansion, and positively selected genes. Our analyses unveiled shrew-specific genomic variants in genes associated with the nervous, metabolic, and auditory systems, which can be linked to unique traits in shrews. Notably, genes suggested to be under convergent evolution in echolocating mammals exhibited accelerated regions in shrews, and pathways linked to putative body size plasticity were detected. These findings provide insight into the evolutionary mechanisms shaping shrew species, shedding light on their adaptation and divergence over time.

## 1. Introduction

Mammals have undergone successive episodes of rapid diversification resulting in the emergence of over 6,000 extant species world-wide (Burgin et al., 2018). Shrews (Soricidae) appeared over 30 million years ago (Churchfield, 1990), and have become one of the most diverse mammalian taxa, having undergone numerous colonization, speciation, and extinction events. (Reumer, 1989). Shrews consist of more than 400 extant species (Burgin et al., 2018) classified into three sub-families; Soricinae (red-toothed shrews), Crocidurinae (white-toothed shrews), and Myosoricinae (African shrews) (Hutterer, 2005). This diversification is also reflected in the diverse karyotypes both between (Schlitter et al., 1999) and within species (White et al., 2019; Wójcik et al., 2002), including sex chromosomes (Sharman, 1956).

Shrews have successfully colonized diverse habitats across the globe (George, 1986, 1988) ranging from tropical forests to grasslands and arid regions (Churchfield, 1990). This has led to a range of adaptations, such as echolocation to navigate their surroundings (Chai et al., 2020; Forsman & Malmquist, 1988; Tomasi, 1979), aquatic diving capabilities to hunt (Mendes-Soares & Rychlik, 2009), venomous saliva for predation (Kita et al., 2004), metabolic shifts (Thomas et al., 2023) and reversible body size changes to survive winter (Lázaro & Dechmann, 2021). Shrews are among the smallest, shortest-lived mammals (Churchfield, 1990) and have an extremely high metabolic rate (Ochocińska & Taylor, 2005) which requires them to consume up to 125% of their body weight in food each day (Churchfield, 1990). These adaptations and overall diversity of shrews make for a unique system to investigate genome evolution in the context of mammalian diversity and evolution.

Approximately 10% of the human genome appears evolutionary constrained across mammals (Christmas et al., 2023). These highly conserved regions are predicted to be functionally important (Bi et al., 2023; Ponting, 2017), but might also experience an increase in nucleotide substitutions, which are referred to as accelerated regions (ARs) (Ferris et al., 2018; Pollard, Salama, King, et al., 2006). Accelerated regions are thought to contribute to species-specific traits (Ferris et al., 2018; Levchenko et al., 2018) and can result from evolutionary forces such as positive selection, GC-biased gene conversion, and relaxed negative selection (Bi et al., 2023; Ferris et al., 2018; Hubisz & Pollard, 2014; Pollard, Salama, King, et al., 2006). Ferris *et al*. (2018) and Tollis *et al*. (2021) identified ARs enriched near immune system and DNA damage response genes in elephants (*Loxodonta*), which are known to show resistance to cancer. Similarly, human ARs have been associated with proteins hypothesized to be important in neurodevelopment (Pollard, Salama, Lambert, et al., 2006 add).

On a larger scale, gene duplication provides new genetic material that can generate novel phenotypes that selection can act upon (Magadum et al., 2013). In bats (*Myotis*), repeated duplications of the protein kinase R (*PKR*) gene have been linked to immunity to viruses (Jacquet et al., 2022). Similarly, gene loss also has the potential to lead to phenotypic evolution and diversity (Helsen et al., 2020; Sharma et al., 2018). The loss of AMP deaminase 3 (*AMPD3*) in sperm whales (*Physeter macrocephalus*) has likely improved O_2_ transport (Sharma et al., 2018). Other variants, such as larger structural rearrangements, also contribute to genomic diversity and evolution (Damas et al., 2021, 2022), highlighting the complex nature of the genomic architecture underlying phenotypic evolution.

Comparative genomics can be used to identify unique and shared genomic features, providing insight on the genetic basis of diversity and patterns involved in species-specific traits. Understanding gene family evolution, including adaptive expansion (i.e., duplication of genes) and adaptive contraction (loss of genes), has increasingly become a focus of molecular evolutionary genetics as the number of fully sequenced genomes has been increasing (Rhie et al., 2021). Here we assembled two *de novo* shrew genomes for the maritime shrew (*Sorex maritimensis*), which is endemic to Canada (Stewart et al., 2002), and the smoky shrew (*S. fumeus*) (Figure S1a). Although restricted to Canada, the maritime shrew is part of the *Sorex* subgenus of *Sorex* shrews that are predominantly found in Eurasia (Fumagalli et al., 1999). The smoky shrew is a member of the subgenus *Otisorex*, members of which are predominantly found in the Nearctic region (George, 1988). We included two previously assembled shrew genomes for the Etruscan shrew (*Suncus etruscus*) and the Eurasian common shrew (*Sorex araneus*) and compared them against 16 other mammal genomes (Figure S1). We aimed to uncover shrew-specific genomic changes associated with their distinctive phenotypes, with the prediction that shared genomic variants among shrew species will be correlated with unique phenotypes and traits shared among these species. We focused on characterizing accelerated regions, gene family duplications, expansion, and contraction events, and positively selected genes in shrew to provide insights into the evolutionary mechanisms driving shrew diversity and adaptation.

## 2. Materials and Methods

### 2.1 Sampling and sequencing

A single smoky shrew (*S. fumeus*) was live captured from Peterborough, Ontario (Animal Care Certificate no. 26234). Heart, liver, brain, and tail tissue were collected immediately after euthanasia and frozen. DNA was extracted from the heart and liver using the *MagAttract HMW DNA Kit* from QIAGEN. Pooled DNA extracts were sent to The Centre for Applied Genomics (TCAG) in Toronto, Ontario, Canada, to sequence 5 HiFi cells on the PacBio Sequel II instrument. Tissue samples were sent to Phase Genomics (Seattle, Washington) to generate a Hi-C library using the Phase Genomics Proximo Animal kit version 4.0.

Genomic sequencing of the maritime shrew was undertaken as part of the CanSeq150 program (https://www.cgen.ca/canseq150-project-list). A single maritime shrew (*S. maritimensis*) was live captured and euthanized following the guidelines of the Canadian Council on Animal Care– see Dawe et al. (2009). Liver tissue was collected immediately and frozen. Genomic DNA was extracted from frozen tissues using a standard proteinase K–phenol–chloroform protocol (Sambrook et al., 1989). DNA and tissues have been stored at -80° and DNA extracts from this maritime shrew were sent to TCAG for sequencing. Short-read (SR) libraries were generated using PCR-free preparation and sequenced on two lanes on the Illumina HiSeqX instrument and with 150 bp paired-end reads. Linked reads (LR) were prepared using the 10x genome library after selecting fragments >15 kb using BluePippin. The LR were sequenced on one Illumina HiSeq X Lane with 150 bp paired-end reads.

### 2.2 Genome assemblies and annotation

We assembled the smoky shrew reference genome following the updated Vertebrate Genome Pipeline (Rhie et al., 2021). The HiFi bam files were filtered using bamtools version 2.5.1 (Barnett et al., 2011) (rq >=0.99) and converted to fastq files using samtools version 1.12 (Danecek et al., 2021) via the *fastq* flag. We trimmed residual HiFi adaptor sequences using cutadapt version 3.4 (Martin, 2011). The primary assembly was constructed using hifiasm version 0.16.1-r375 (Cheng et al., 2021, 2022) allowing for the integration of the Hi-C reads. We used the default level of purge duplication of 3 for non-trio assembly. We ran the hifiasm python module on 32 threads with 228 GB of RAM. We used Merqury version 1.3 (Rhie et al., 2020) with meryl from Canu version 2.2 (Koren et al., 2017) to assess genome assembly quality via *k*-mer copy number analysis.

For the maritime shrew genome, we used BBMap version 35.8 (Bushnell, 2022) to remove adapters and trim low-quality data from the fastq files. Kmergenie version 1.7048 (Chikhi & Medvedev, 2014) was used to find the kmer size for downstream analysis. The genome was assembled using a tiered approach: (1) using only the SR data with w2rap-contigger with -K set to 144 (Clavijo, 2021) and (2) with only the 10x LR data using Supernova version 2.1.1 (Weisenfeld et al., 2018) using the --maxreads filter set to “all”. The pseudohap2 style was selected for the assembly output. Following these assemblies, both SR and LR assembly versions were merged via quickmerge version 0.3 (Chakraborty et al., 2016) with the LR as the backbone and the default parameters. Scaff10x version 5.0 (Ning et al., n.d.) was used to further polish the maritime shrew genome.

Nuclear genome completeness was assessed for both assemblies using Benchmarking universal single-copy orthologues (BUSCO) version 3.0.2 (Simão et al., 2015) by comparing the genomes to highly conserved genes in mammals. The smoky shrew mitochondrial genome was assembled using the PacBio subreads and the mitochondrial genome of a previous smoky shrew assembly (NCBI accession: GCA_026122425.1) (Cossette et al., 2023) as a backbone with MitoHifi version 3.2 (Allio et al., 2020; Uliano-Silva et al., 2023). The mitochondrial genome for the maritime shrew was assembled using MitoZ (Meng et al., 2019) with the function “mitoz all” on the fastq reads and parameters --genetic_code 2 --clade Chordata --kmers_megahit 59 63 79 99 119 141 --requiring_taxa Chordata and was manually curated.

The smoky shrew genome annotation was generated by NCBI using their eukaryotic genome annotation pipeline the integrated RNAseq data (SRA RNA-Seq accession: SRX20204431, SRX20204430) from Cossette *et al*. (2023). We generated the maritime shrew annotation in-house via GenSAS version 6.0 (Humann et al., 2019) as it was too fragmented for the NCBI pipeline. Repeat regions were identified using RepeatModeler version 2.0.1 (Smit & Hubley, 2023b) and RepeatMasker version 4.1.1 (Smit & Hubley, 2023a). Here, we had to use the smoky shrew liver and heart RNA reads from Cossette *et al*. (2023) and mapped them to the genome using HISAT2 version 2.2.1 (Kim et al., 2019). The resulting BAM files were used by Augustus version S3.4.0 (Stanke et al., 2006) for gene prediction. The NCBI refseq vertebrate-mammalian protein database available on GenSAS was aligned to the genome using DIAMOND version 2.0.11 (Buchfink et al., 2015). Augustus and DIAMOND were also run using the common shrew, mSorAra2.pri (GenBank accession: GCF_027595985.1), protein fasta file. EVidenceModeler version 1.1.1 (Haas et al., 2008) was used to generate a consensus gene set using the Augustus (1x weight) and DIAMOND (5x weight) outputs. Gene function was assigned to our gene consensus model with DIAMOND and InterProScan version 5.53-87.0 (Jones et al., 2014)

### 2.3 Detection of ARs in shrew genomes

We generated a multiple alignment file (MAF) consisting of 20 mammalian genome assemblies to identify ARs in the four shrew species. We downloaded the pairwise syntenic net alignment files for the 16 mammals, including the common shrew, against the human (hg38) genome (Table S1) from the UCSC database (Kent et al., 2002). Using LASTZ version 1.04.03 (Harris, 2007) we generated pairwise alignments against the hg38 genome for the smoky, maritime, and Etruscan shrew genomes. The LASTZ run parameters were set to K=3000, L=3000, Y=9400, E=30, H=2000, and O=400 based on UCSC’s hg38 100-way conservation parameters for their common shrew alignment. The alignments for each species were chained with the chainMinScore=3000 and linearGap=medium options and subsequently netted using kentutils tools version 401 (UCSC Genome Browser, 2020). Our resulting pairwise alignments for each shrew species were then aligned to the other mammal pairwise alignments to create one MAF using the roast function from MULTIZ version 11.2 (Blanchette et al., 2004) and the tree topology in Figure S1b. The required tree was inferred by Orthofinder (see section 2.4) and supported by Murphy *et al*. (2001) and Meredith *et al*. (2011).

The MAF was filtered to only keep alignments to the 22 main autosomal chromosomes of the hg38 genome. We used the msa_view function from the PHAST package version 1.5 (Hubisz et al., 2011) to extract 4-fold degenerate (4D) sites from the non-shrew species in our alignments and based on the human annotations (NCBI accession: GCA_000001405.15) from which we only kept coding sequences. The output was used to estimate a nonconserved phylogenetic model using phyloFit with the substitution model REV and the EM algorithm option. The PhastCons function from RPHAST version 1.6.11 (Hubisz et al., 2011) was used to identify conserved regions in the non-shrew species with the parameters set to expected.length=45, target.coverage=0.3, rho=0.31 and viterbi=TRUE. We filtered the predicted conserved regions to sites that aligned to all the shrew species and at least 18 species in total. We regularized the length of the conserved regions to 50 bp regions to simplify the likelihood ratio tests. PhyloP was run using the ACC mode to identify accelerated regions in each of our shrew species. Non-parametric simulations were run to calculate empirical p-values. Statistically significant ARs were defined with a false discovery rate threshold of 5%. Genes overlapping these ARs, were identified using bedtools version 2.30.0 (Quinlan & Hall, 2010) and the hg38 genome annotations.

### 2.4 Gene family size evolution and non-synonymous to synonymous rate ratio

We identified orthogroups between proteins sequences from all 20 mammal genomes (Table S1, Figure S1) to identify gene duplication events. We downloaded the protein files from NCBI RefSeq for each species that included the smoky shrew (Table S1), except the maritime shrew for which we used our GenSAS annotation. We further filtered to keep the longest transcript variant per gene in each species’ file to minimize run time and increase accuracy. OrthoFinder version 2.5.4 (Emms & Kelly, 2019) with the parameters -M msa, -S diamond, -A mafft, -z, -T fasttree was used to obtain orthogroups and identify duplicated genes. We then used CAFÉ version 4.2 (De Bie et al., 2006) with a multi-lambda model to analyze the evolution of orthogroup sizes and used an error estimation model to account for genome assembly errors.

We used an adaptive branch site random effects likelihood (aBSREL) model (Smith et al., 2015) with the HyPhy package version 2.5.49 (Pond et al., 2005) to identify possible episodic diversifying selection in shrews by calculating the ratio of non-synonymous to synonymous substitutions (dN/dS). The analysis was performed on the single-copy genes identified by Orthofinder. Multiple sequence alignment for each protein were converted to sequence codon alignments using PAL2NAL version 14.1 (Suyama et al., 2006) to input in aBSREL using the coding sequence available on RefSeq (Table S1). We ran the aBSREL analysis twice to estimate dN/dS, first (i) with the branch leading to the common ancestor of the shrews and all its descendent branches selected as the foreground; and (ii) with no foreground branches specification and all branches are tested for positive selection. An R script (Jarva, 2023) was used to parse and extract data from the nested json file outputs.

### 2.5 Gene pathway analysis

Gene and protein summaries were manually retrieved from UniProt (Bateman et al., 2023) and NCBI (National Center for Biotechnology Information (NCBI), 1988). Genes that were associated with ARs, duplications events, significant family size changes, and positive selection in shrews were linked to the corresponding human ortholog gene ID when possible, to input in the Database for Annotation, Visualization and Integrated Discovery (DAVID) (Huang et al., 2009; Sherman et al., 2022), to identify gene pathways for each analysis.

## 3. Results

### 3.1 Genome assembly and annotations

For the smoky shrew assembly (Genbank assembly accession: GCA_029834395.2), the five cells produced 115,806,042,303 bp of HiFi data, or ∼43X coverage from the estimated 2.7 GB genome (Cossette et al., 2023). The mean length of the HiFi reads was 14,652 bp. We generated 368,278,690 paired Hi-C reads. Using the Hi-C reads for scaffolding the assembly, the final assembly smoky shrew assembly was 2.87 GB with an N50 of 42.40 MB and over 99% of the 499 scaffolds being >50 KB long (Table 1). Approximately 51.9% of the genome was made of repetitive sequences (Table 2). The BUSCO assessment was 93.4 % complete, of which 91.8 % were single-copy. Distribution plots showed a high-percentage of single copy k-mers in the final assembly. We generated 890,398,361 raw 150 bp paired-end reads (∼110X coverage) and 96,242,024,700 bp of 10x reads. The maritime shrew genome was 2.43 GB (GenBank assembly accession: GCA_030324115.1) and comprised of 91,327 scaffolds with an N50 of 84.5 KB. 69.8 % of the scaffolds were over 50 KB long and 41.7 % of the genome was classified as repetitive sequences. Out of the BUSCO orthologs from the mammalian data set 72.9 % were complete with 70.6 % being single-copy.

**Table 1.**
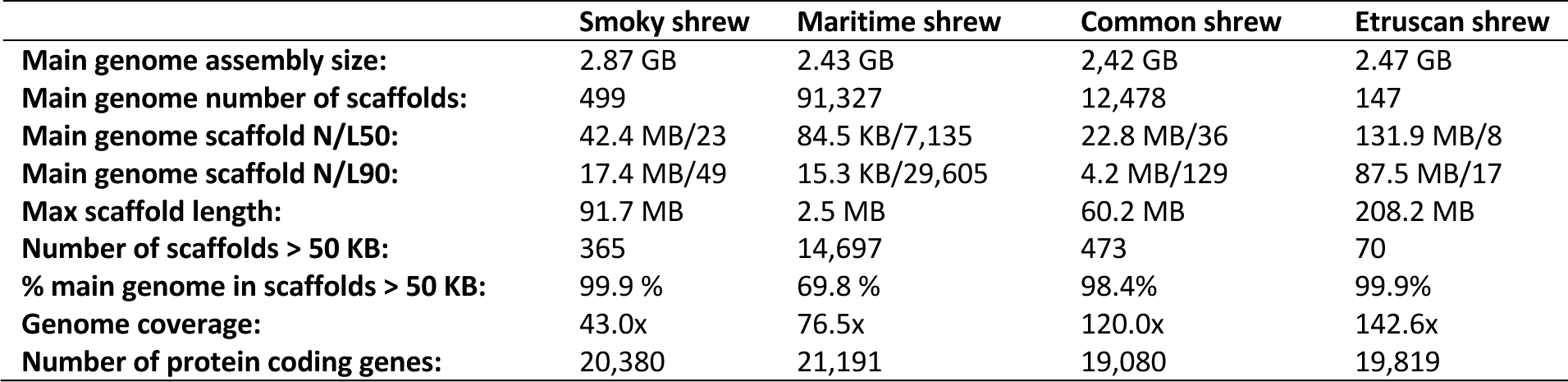
Genome assembly statistics for the smoky shrew genome generated from PacBio HiFi long-reads and Hi-C scaffolding, maritime shrew genome generated from 10X long-reads and short-reads, and the common shrew and Etruscan shrew genomes downloaded from NCBI.

**Table 2.** Summary of repeats in the smoky and maritime shrew genomes in length occupied (bp) and percentage of the genome.

|  | Smoky shrew | Maritime shrew |
| --- | --- | --- |
| SINE | 182,692,788 (6.36 %) | 141,858,027 (6.90 %) |
| LINE | 614,478,406 (21.38 %) | 366,060,918 (17.81 %) |
| LTR elements | 41,000,372 (1.43 %) | 30,015,478 (1.46 %) |
| DNA transposons | 10,884,988 (0.38 %) | 6,592,309 (0.32 %) |
| Unclassified | 500,269,535 (17.41 %) | 198,021,546 (9.63 %) |
| Rolling-circles | 686,527,20 (2.39 %) | 58,004,316 (2.82 %) |
| Small RNA | 20,490,319 (0.71 %) | 18,114,174 (0.88 %) |
| Satellites | 2,682,795 (0.09 %) | 2,057,146 (0.10 %) |
| Simple repeats | 39,822,407 (1.39 %) | 29,813,949 (1.45 %) |
| Low complexity | 9,066,276 (0.32 %) | 5,801,032 (0.28 %) |
| Total | 1,421,387,886 (51.86 %) | 856,338,895 (41.65 %) |

A total of 20,380 protein coding genes were identified in the smoky shrew genome and 21,191 in the maritime shrew genome. The assembled maritime and smoky shrew mitochondrial genomes were 16,979 bp and 17,051bp long respectively; all 37 mitochondrial genes were present in both assemblies.

### 3.2 Accelerated regions

We identified 598,180 conserved regions (Table S2; ∼1 % of the genome) in our alignments, comparable to other mammalian studies (Ferris et al., 2018; Tollis et al., 2021). From these regions, 307,989, ∼ 51% of conserved regions, were in exons. We found 2,643 common shrew ARs in 1,627 genes, 21,581 maritime shrew ARs in 4,528 genes, 4,272 smoky shrew ARs in 2,257 genes, and 21,354 Etruscan shrew ARs in 6,787 genes (Table 3, S2). Soricidae shared 35 ARs and the *Sorex* species shared 404 ARs (Figure 1a). Notable ARs were found in genes related to the nervous system such as the growth associated protein 43 (*GAP43*), fibroblast growth factor receptor 1 (*CPNE6*), class III β-tubulin (*TUBB3*) (Table 4). Also, genes involved in metabolic pathways such as the adiponectin receptor 1 (*ADIPOR1*) and glyceraldehyde-3-phosphate dehydrogenase (*GAPDH*) had shared ARs in *Sorex* and Soricidae respectively (Table 4). The common shrew and maritime shrew genomes shared ARs in the cadherin related 23 (*CDH23*) and otoferlin (*OTOF*) genes, which are associated with echolocation capabilities in mammals (Shen et al., 2012; Table 4). The pathway analyses revealed genes associated with nervous system development, olfactory receptors, and insulin and energy metabolism pathways (Figure 1b).

**Figure 1.**
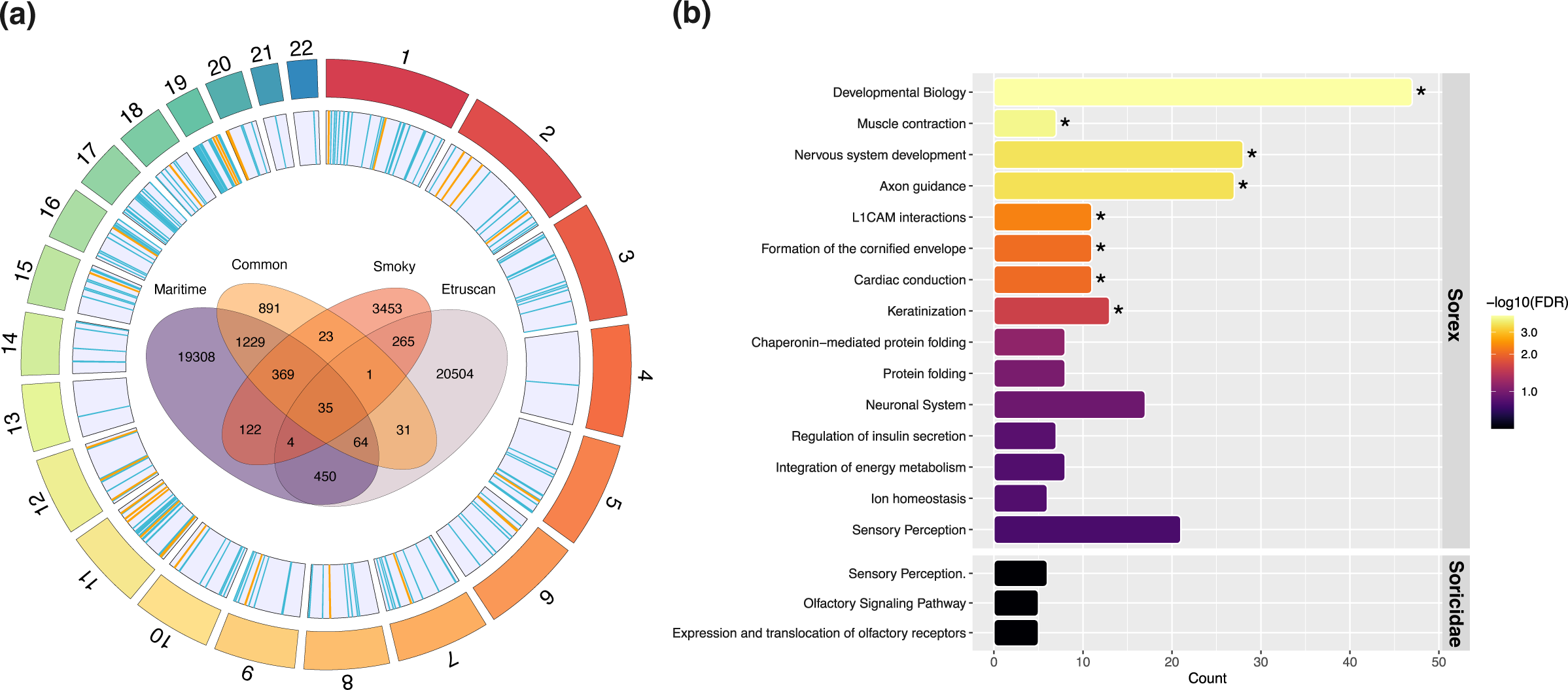
Accelerated region (AR) analysis result **(a)** Location of ARs shared between Sorex species (n=404) in blue and Soricidae (n=35) in yellow mapped to the human (hg38) genome. Number of overlapping ARs between shrew species. **(b)** Top Reactome pathways for overlapping ARs in Sorex and Soricidae. Asterisk represents significant pathways (5% FDR).

**Table 3.** Summary statistics from each analysis per shrew species/node.

|  | Smoky shrew | Maritime shrew | Common shrew | Etruscan shrew | Sorex | Soricidae |
| --- | --- | --- | --- | --- | --- | --- |
| # accelerated regions | 4,272 | 21,581 | 2,643 | 21,354 | 404 | 35 |
| # gene duplications | 2,328 | 2,658 | 2,722 | 2,306 | 62 | 42 |
| # rapidly +/- orthogroups | +85/-8 | +54/-503 | +407/-1 | +128/-36 | +62/-5 | +52/-18 |
| # genes under selection | 31 | 97 | 9 | 116 | 62 | 46 |

**Table 4.**
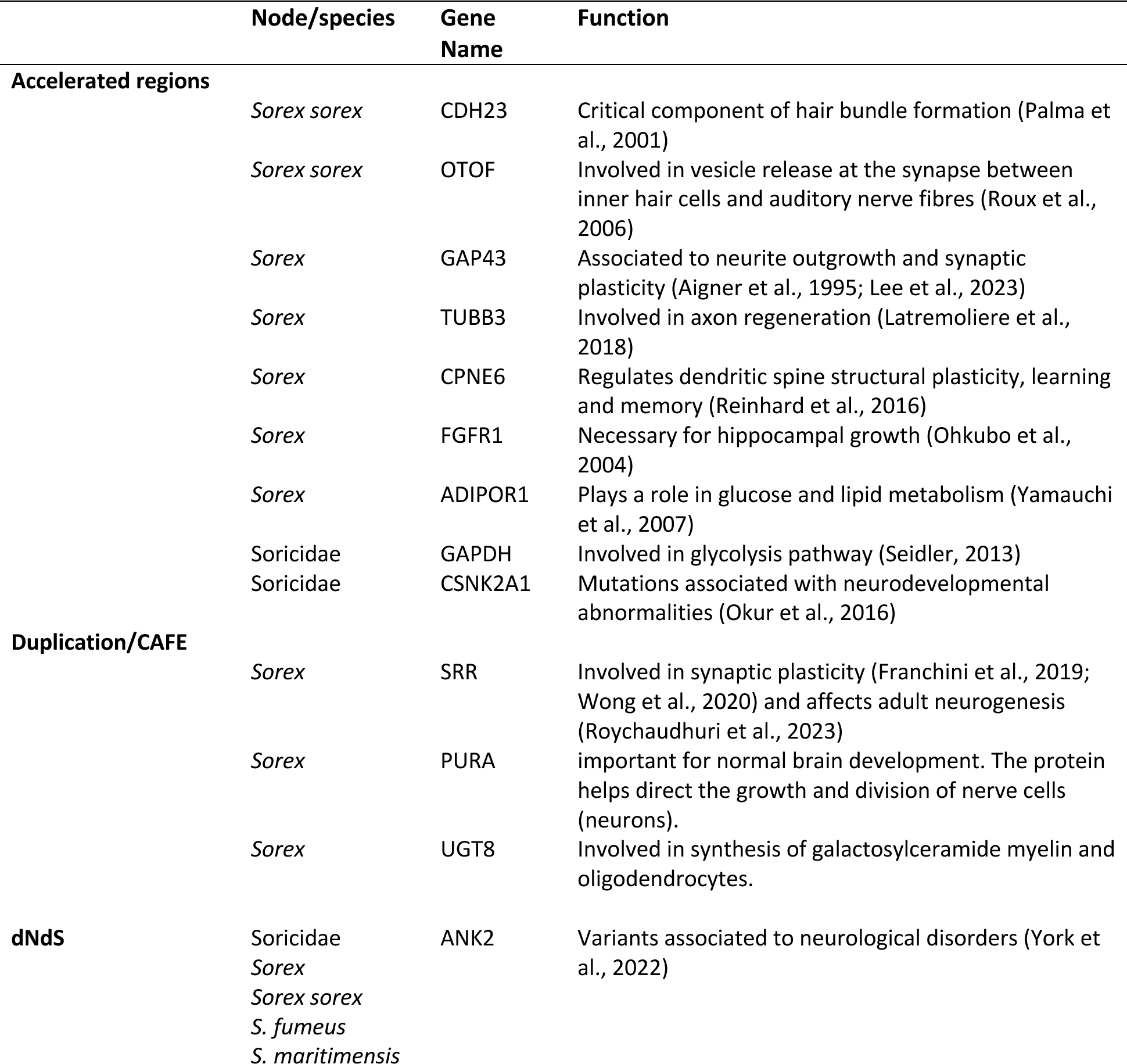
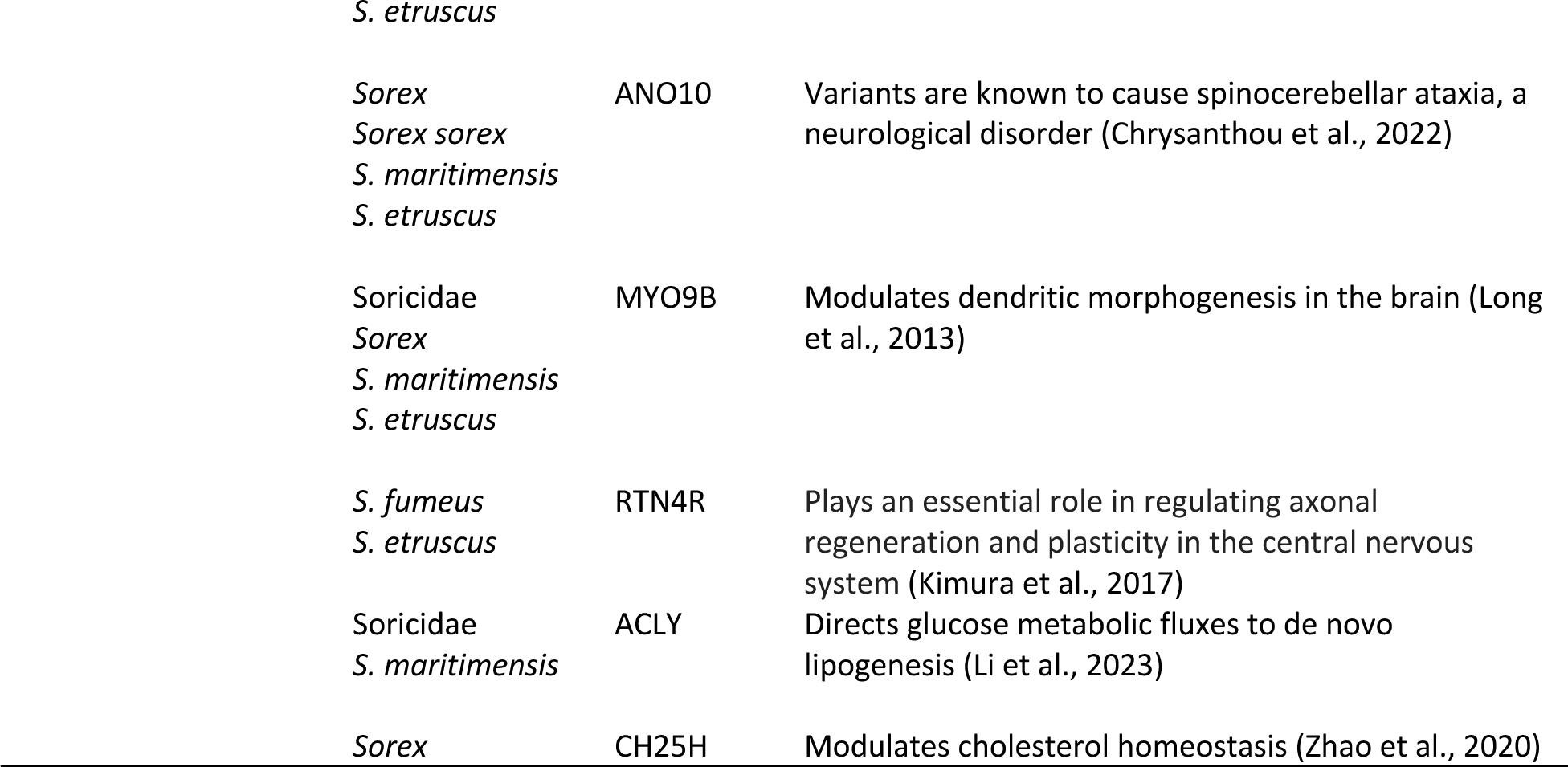
Relevant genes in shrew species/nodes identified in each analysis based on the human hg38 annotations (NCBI accession: GCA_000001405.15). References in supplemental Table S3.

### 3.3 Gene family size evolution

Orthofinder assigned 476,077 (97.0% of total) genes to 20,489 orthogroups. Of these, 7,954 orthogroups included at least one ortholog from each of the 20 species and 1,810 orthogroups were species-specific. A total of 533 genes were identified as single-copy genes across all 20 mammals. The maritime shrew had 15,755 (74% of genes) genes assigned to orthogroups (Figure 2a) and had the most species-specific orthogroups (Figure 2b). The three other shrew species’ assemblies had on average 37 unique gene families each (Figure 2b). Orthofinder further identified 2,722 gene duplication events in the common shrew, 2,658 in the maritime shrew, 2,328 in the smoky shrew and 2,306 in the Etruscan shrew (Figure 2c, Table 3). There were 42 gene duplication events shared between all shrew species and 62 between the S*orex* genus (Figure 2c, Table 3).

**Figure 2.**
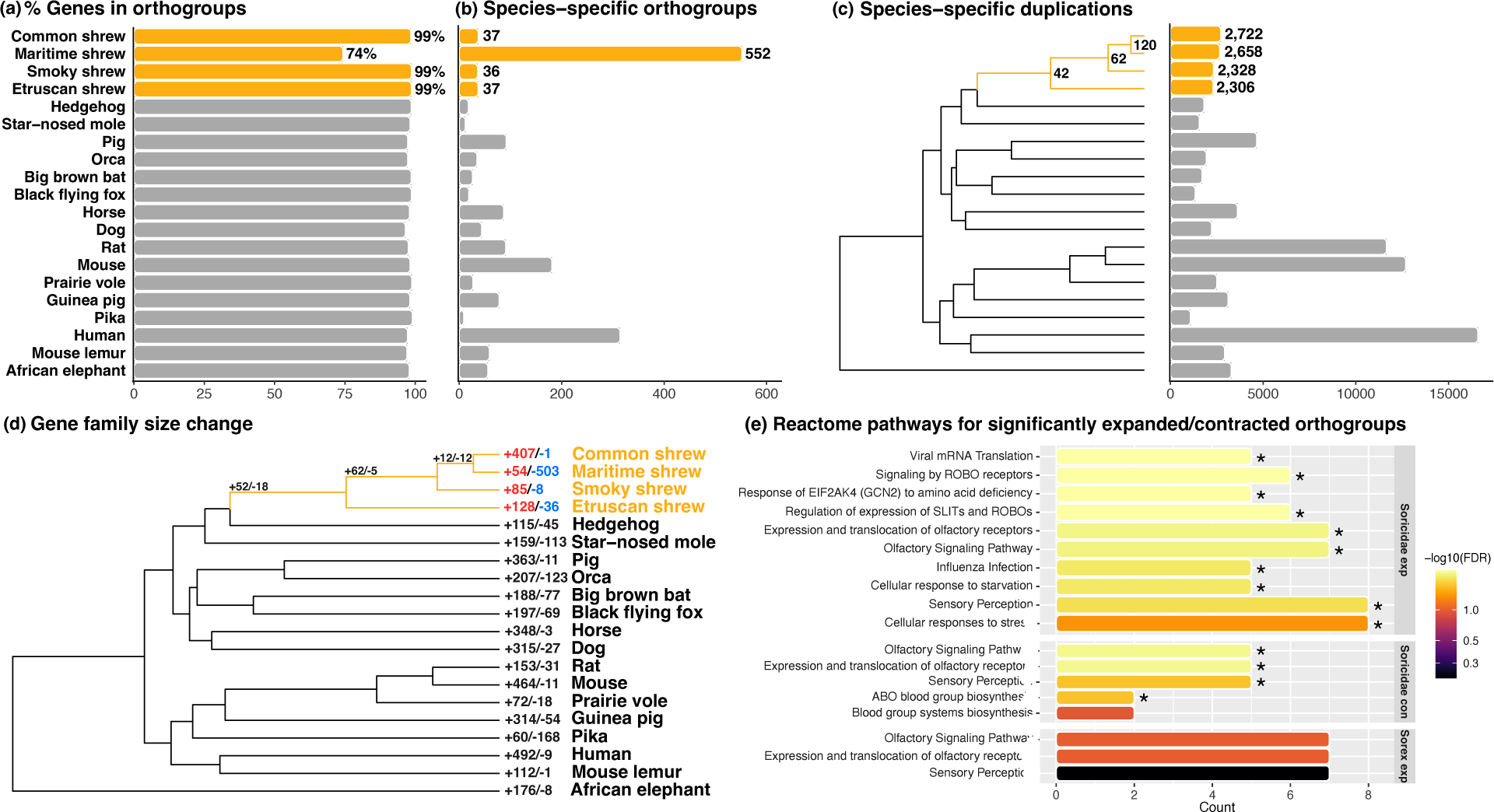
OrthoFinder and CAFE analyses results **(a)** Percentage of genes from each species assigned to orthogroups. **(b)** Number of species-specific orthogroups for each species. **(c)** Number of species-specific duplication events. Number of duplication events shared by all descendants of the shrew nodes are also represented. **(d)** Number of significantly expanded/contracted orthogroups across 20 mammalian species. **(e)** Top Reactome pathways for the significantly expanded/contracted orthogroups shared between Sorex and Soricidae species. Asterisk represents significant pathways (5% FDR).

The number of gene families undergoing expansion ranged between 904 to 1,476 in shrews with up to 407 rapidly expanding families in the common shrew (Figure 2d). The maritime shrew appeared to have undergone more contractions (n=7,285) than any other shrew species which ranged between 629 to 1,598 contractions (Figure 2d, Table 3). The orthogroup associated with the serine racemase (SRR) gene, which is involved in neurogenesis (Roychaudhuri et al., 2023), appeared to be rapidly expanding in the *Sorex* shrews (Table 4). Other significantly expanding gene families in all Soricidae were associated with nervous system development genes and cellular response to stress and starvation, olfactory receptors, and the immune system (Figure 2e). The gene families undergoing significant contractions shared between species were related to olfactory receptors and blood group system pathways (Figure 2e).

### 3.4 Episodic diversifying selection in shrews

We calculated the ratio of non-synonymous to synonymous substitutions (dN/dS) using aBSREL to identify positive selection. Out of the 533 single-copy genes tested, nine were under selection in the common shrew, 97 in the maritime shrew, 32 in the smoky shrew and 116 in the Etruscan shrew when the Soricidae common ancestor and all descendent branches were selected as the foreground branches (Figure 3a, Table 3). A total of 62 genes were under positive selection in the Soricidae node, and 46 were selected for in the *Sorex* node (Figure 3a, Table 3). In all shrew nodes and species, except for the common shrew, the ankyrin 2 (*ANK2*) gene, involved in making proteins found in the brain and heart was under positive selection (Figure 3b, Table 4). Other genes under selection included anoctamin 10 (*ANO10*), myosin IXB (*MYO9B*), and reticulon 4 receptor (*RTN4R*) which are all associated with the nervous system (Table 4). Top enriched pathways for positively selected genes were related to protein metabolism and various GTPases cycles (Figure 3c).

**Figure 3.**
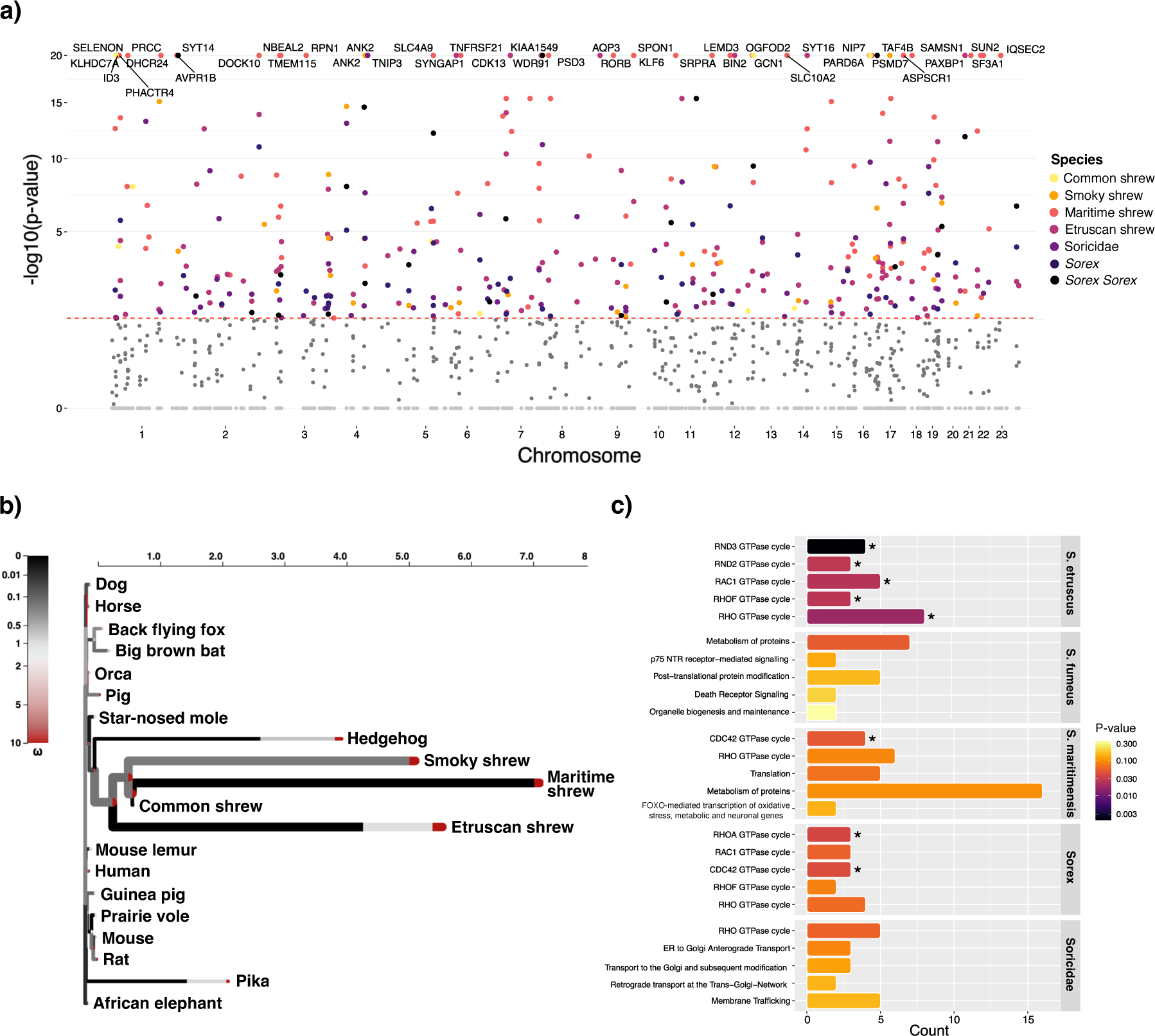
aBSREL (dN/dS) analysis results **(a)** Manhattan plot of Bonferroni-Holm corrected p-values used as evidence for selection for all 533 single-copy genes from the foreground run. Dark grey points represent non-significant p-values shrew nodes and species. Light grey points represent all other mammal nodes and species. The red dotted line represents the 0.05 significance threshold. Top shrew nodes and species are indicated by the respective gene name. **(b)** HyPhy Vision plot for the different omega (ω) rate distributions in each branch for the ANK2 gene. **(c)** Top Reactome pathways for the positively selected genes from the dN/dS analysis in shrew nodes and species. Asterisk represents significant pathways (p-value < 0.05).

## 4. Discussion

Over the course of mammalian evolution, species have adapted to diverse ecological niches, evolving an array of specialized traits and behaviors. Shrews, in particular, have evolved unique phenotypes and abilities tailored to their various environments, including venomous saliva (Kita et al., 2004), echolocation (Chai et al., 2020; Forsman & Malmquist, 1988; Tomasi, 1979), hunting underwater (Mendes-Soares & Rychlik, 2009), and reversible body size changes (Lázaro & Dechmann, 2021). Soricidae are also known to have one of the shortest lifespans and fastest mass-specific metabolic rate of all mammals (Ochocińska & Taylor, 2005). Here we assembled two *de novo* shrew genomes and conducted comparative genomic analyses between four shrews and 16 other mammal species to test our prediction that shared genomic variants among shrew species are likely associated with unique phenotypes and abilities specific to shrews. We identified accelerated regions, gene family expansion and contractions, as well as genes undergoing positive selection in Soricidae.

### 4.1 Genome evolution underling phenotypic changes in shrews

Shrews have poor vision which is mostly used to detect light intensity; in contrast, their olfactory, tactile, and acoustic senses are well developed (Churchfield, 1990). Certain Nearctic shrew species, and the common shrew, have even developed echolocating abilities to navigate their surroundings (Churchfield, 1990; Forsman & Malmquist, 1988; Gould, 1969). We found shared ARs in the coding region of the *CDH23* gene and 3’UTR region of the *OTOF* gene in the common and maritime shrews (Table 4). Both genes are involved in the auditory system and more specifically echolocation in mammals (Shen et al., 2012). Chai *et al*. (2020), found evidence of convergent evolution between the common shrew and other echolocating mammals for both these genes. The common shrew is the only species including in our analyses confirmed to echolocate, but our findings indicate the potential for such ability in the maritime shrew, as it is the most closely related species and shares ARs with the common shrew for the *CDH23* and *OTOF* genes. Echolocation is likely more widespread among shrews (i.e *Blarina*) (Gleason et al., 2023), including other *Sorex* (Buchler, 1976), thus detection in the maritime shrew is not unexpected and this might be a *Sorex*-wide trait.

Survival in cold climates involves species adopting strategies such as migration, hibernation, or entering a state of torpor. Shews do not hibernate nor migrate long distances (Churchfield, 1990). Their high metabolic rate prevents them from building up sufficient fat storage (Churchfield, 1990); therefore, to survive through the winter, some shrew species have evolved a way to lower their energetical demand by undergoing reversible seasonal changes in body size and mass, known as Dehnel’s phenomenon (Lázaro & Dechmann, 2021). During the winter, brain mass can shrink around 20% in the common shrew and regrow up to 17% for the summer (Lázaro et al., 2017). This has also been shown in the Etruscan shrew (Ray et al., 2020), and other *Sorex* shrews (e.g. *Sorex minutus*) (Bartkowska et al., 2008; Lázaro & Dechmann, 2021). It remains uncertain if the maritime shrew and smoky shrew exhibit Dehnel’s phenomenon, however both species belong to the *Sorex* genus and inhabit Northern climates, akin to other shrew species known to undergo these size changes.

We observed hundreds of ARs in shrews associated with nervous system development and axon guidance pathways (Figure 1b), including specific genes such as *TUBB3*, *GAP43* and *FGFR1* (Table 4). In all *Sorex* species, ARs were found in the 5’UTR regions of the *TUBB3* an *GAP43* genes. *TUBB3* plays a role in the regeneration of axons (Latremoliere et al., 2018), while *GAP43* is involved in neurite outgrowth and synaptic plasticity in the hippocampus (Aigner et al., 1995; Lee et al., 2023), a region of the brain known to shrink during the winter and regrow in the spring in the common shrew (Lázaro et al., 2018). Furthermore, ARs were identified in the coding region of the *FGFR1* gene, which is also involved in hippocampal growth (Ohkubo et al., 2004). These findings were further supported by the gene family expansions. Gene families associated with the *SRR*, *PURA* and *UGT8* genes, which are all involved with the brain and the nervous system (Table 4), were undergoing rapid expansion in all *Sorex* species. And in all Soricidae species, significantly expanding gene families were associated with SLITs and ROBOs pathways (Figure 2e) which are involved in neocortical development, a region responsible for sight and hearing (Gonda et al., 2020). The pathway analysis further revealed that positively selected genes in shrew were related to Rho GTPase pathways (Figure 3). Rho GTPases, such as RhoA, Cdc42, Rac1 are involved in neuronal development, neurodegeneration, and synaptic plasticity (Stankiewicz & Linseman, 2014; Zhang et al., 2021). Finding such genes and pathways associated with the brain and nervous system consistently in all analyses and shared between all species we hypothesize is indicative of the genetic mechanisms involved in seasonal brain size plasticity.

Shrews might also undergo a metabolic shift from lipid to glucose as <u>a</u> fuel source to survive the winter. Work by Thomas *et al*. (2023) found evidence that lipid metabolite concentration decreased throughout the winter in the common shrew, possibly promoting carbohydrate metabolism during the harsh winter months. Thomas *et al*. (2023) found differentially expressed genes and pathways associated with insulin and cholesterol between shrew brain and liver samples across seasons. When consumed, carbohydrates breakdown into glucose, and insulin then regulates if it is used as source of energy or stored (Dube et al., 2013). We identified genetic variants in the AR and dN/dS analyses associated with Rho GTPase pathways in the accelerated region analysis (Figure S1, 3). Rho GTPase are also involved in metabolic homeostasis more specifically glucose metabolism (Møller et al., 2019). Furthermore, an accelerated region was identified in the 3’UTR region of the *ADIPOR1* gene in *Sorex* (Table 4). *ADIPOR1* plays a crucial role in the regulation of glucose and lipid metabolism (Yamauchi et al., 2007). All shrew species shared an accelerated region in the *GAPDH* coding region, which is involved in glycolysis, the process that breaks down glucose into energy (Table 4). Furthermore, significantly expanding gene families were associated with cellular response to stress and starvation (Figure 2e). These findings highlight the relationship between genomic variants unique to shrews and their distinctively fast metabolism and wintering strategy and offers candidate genes for further hypothesis testing (Table 4).

### 4.2 Avoiding bias from assembly and annotation differences

The divergent assembly strategies and genome quality statistics (Table 1) presented us with a natural study to compare the impact of reference genome that warrants comment. Assembly inconsistencies between genomes can lead to overestimation of genomic differences between taxa when conducting comparative genomic analyses (Denton et al., 2014). Our shrew assemblies consisted of three highly contiguous genomes (Etruscan, smoky and common shrew), and the maritime shrew being the most fragmented. In the accelerated region analysis, the maritime shrew and Etruscan shrew assemblies revealed a high number of ARs; however, we found no clear pattern between number of ARs and genome assembly quality or contiguity (Figure S4). For good measure, we repeated the analysis to identify ARs in the African elephant (*Loxodonta africana*) and big brown bat (*Eptesicus fuscus*) and compared these values to the study done by Ferris *et al*. (2018). Our results were consistent with their work, with a significant lower number of ARs in the African elephant (n=1904) compared to the bat (n=20,939; Table S2). Thus, the AR analysis appears to not be impacted by highly fragmented assemblies like the maritime shrew, likely due to the bioinformatic approach of mapping short fragments.

The maritime shrew had less genes assigned to orthogroups and more species-specific orthogroups compared to all other shrews. Here, this is likely due to the maritime shrew’s genome contiguity and not having a standardized RefSeq annotation like all other species. Depending on the sequencing and assembly method, *de novo* genome assemblies can be fragmented resulting in inaccurate structure and number of predicted genes (Denton et al., 2014). However, despite detecting considerably more orthogroups in the maritime shrew, the number of duplicated regions and genes under positive selection did not deviate significantly from the Etruscan shrew (Table 3), suggesting that none of our metrics were dramatical impacted by fragmentation. Still, to avoid biases due to assembly and annotation differences between species, we focused on overall trends found in Soricidae and *Sorex* instead of species-specific trends. This minimizes overestimation of ARs, gene duplication event or gene family expansion/contraction and positively selected genes identified in shrews. Overall, our study found shrew-specific genomic variants in genes associated with the nervous, auditory, metabolic, and olfactory systems, most of which can be linked to unique phenotypes or traits in shrews.

## Data and Resource Availability

Smoky shrew genome and raw sequence reads deposited in GenBank/NCBI under project: PRJNA826195

Smoky shrew genome and raw sequence reads deposited in GenBank/NCBI under project: PRJNA956518

Scripts are uploaded on GitLab: https://gitlab.com/WiDGeT_TrentU/graduate_theses/-/tree/master/Cossette?ref_type=heads

## Author Contributions

MLC, ABAS and DTS conceived the study and collected the samples. MLC and DTS performed the molecular laboratory work. MLC performed the bioinformatic analyses with contribution from ABAS for the smoky shrew genome assembly. MLC and ABAS wrote the manuscript and DTS reviewed it.

## Acknowledgments

This study was supported by CanSeq150 (CGEn) (ABAS, DTS and MLC); Ontario Graduate Scholarship (MLC); Natural Sciences and Engineering Research Council of Canada (NSERC) Alexander Graham Bell Canada Graduate Scholarship-Master’s (MLC), Canadian Graduate Scholarship-Doctoral (MLC), and NSERC Discovery grants (ABAS grant number: RGPIN-2017-03934 and DTS grant number: RGPIN/217175); French American Charitable Trust Scholarship (MLC); Edwin William Curtin & Irene Elizabeth Curtin Graduate Scholarships (MLC); Ontario Graduate Scholarship (MLC); Compute Canada Resources for Research Groups (ABAS grant number: RRG gme-665-ab); Canadian Foundation for Innovation: John R. Evans Leaders Fund (ABAS); Ontario Early Researcher Awards (ABAS grant number: #36905). We would like to thank The Centre for Applied Genomics in Toronto, Ontario, Canada and Phase Genomics Seattle for their expertise and insight with sequencing. We would like to acknowledge that the work for this study was carried out on the traditional territory of the Mississauga (Michi Saagiig) Anishnaabeg and the Mi’kmaq People. We are grateful to have had the opportunity to work on this land and thank the First Peoples for their care, stewardship, and teaching.

