## Supplemental 1 for "Comparative genomics of the world’s smallest mammals reveals links to echolocation, metabolism, and body size plasticity"

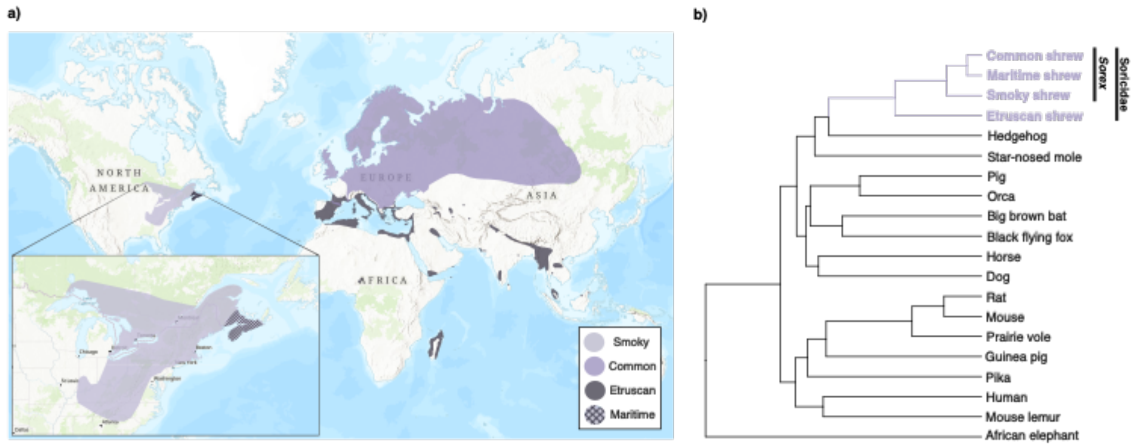


***Figure S1**. **a)** Shrew species' range according to the IUCN (ref). **b)** Tree topology used for the analyses.*

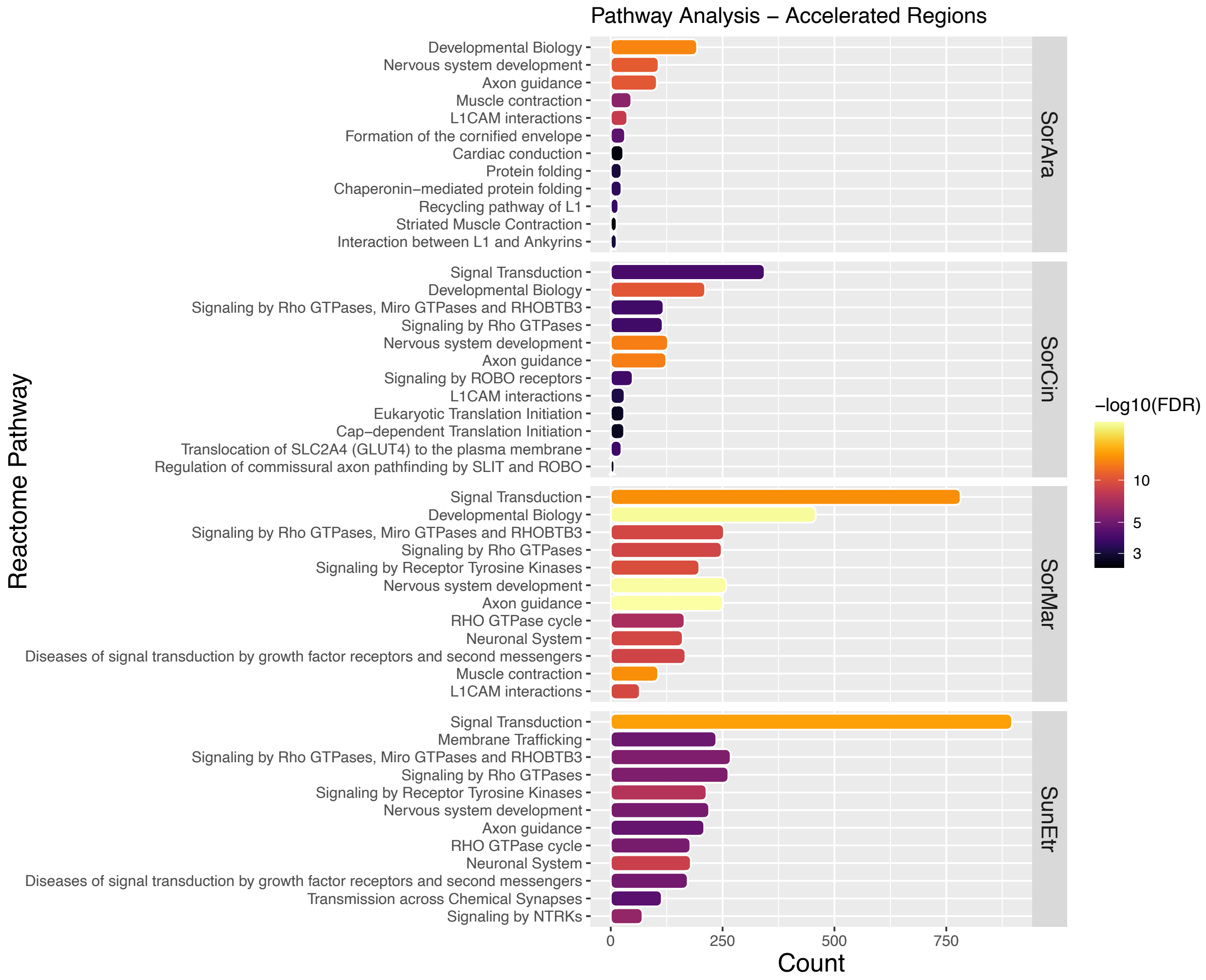


***Figure S3**. Top Reactome pathways for ARs found in each shrew species. All pathways are significant pathways (5% FDR).*

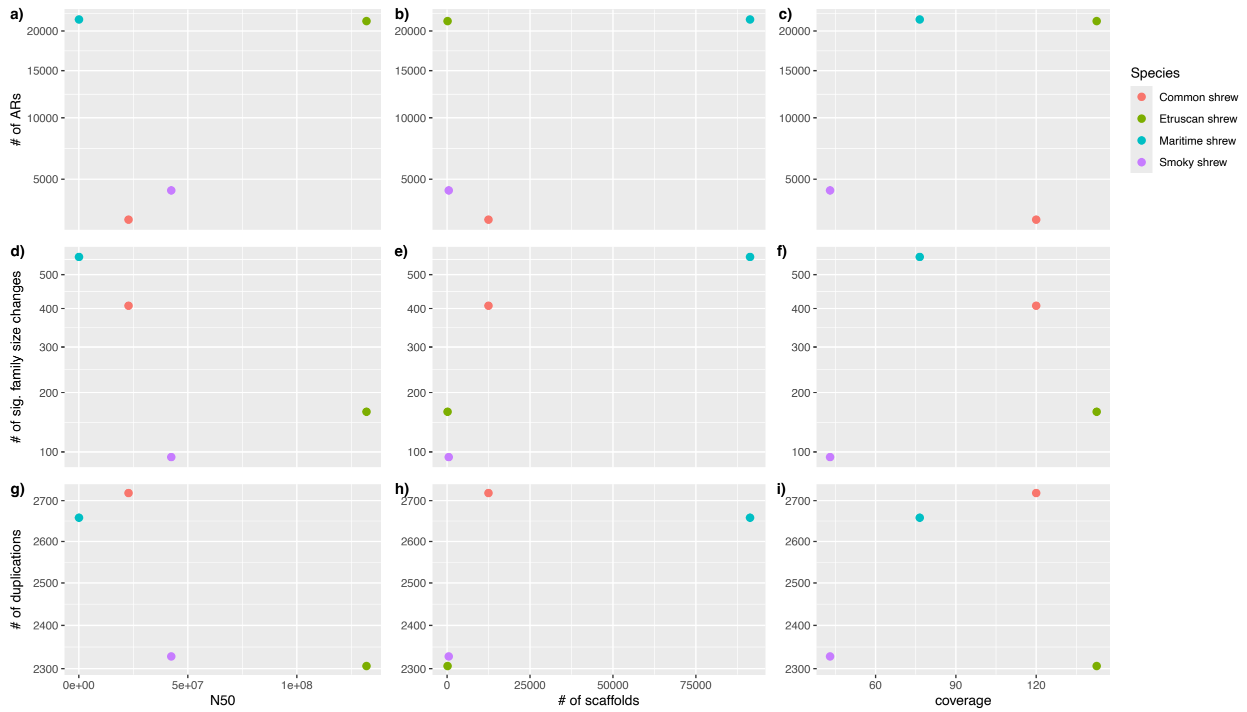


***Figure S4****. Analysis results versus genome assembly statistics for each shrew species.* ***a)*** *Number of ARs versus N50,* ***b)*** *number of scaffolds, and* ***c)*** *coverage.* ***d)*** *Number of gene families undergoing significant size change versus N50,* ***e)*** *number of scaffolds, and* ***f)*** *coverage.* ***g)*** *Number of duplications versus N50,* ***h)*** *number of scaffolds, and* ***i)*** *coverage.*

***Table S1.*** *Species name, genome version, and NCBI RefSeq accession number of the protein.faa and cds_from_genomic.fna files downloaded and used for the OrthoFinder and aBSREL analyses respectively. The synNet.maf files used for each species for the accelerated region analysis matched the genome versions listed here. *There is no RefSeq annotation file for the maritime shrew. We used the maritime shrew protein.faa file from our GenSAS annotation pipeline (available on gitlab).*

| **Species** | **Genome version** | **NCBI RefSeq accession** |
|---|---|---|
| Black flying fox (*Pteropus alecto*) | ASM32557v1 | GCF_000325575.1 |
| Dog (*Canis lupus familiaris*) | CanFam6 | GCF_000002285.5 |
| Domestic guinea pig (*Cavia porcellus*) | Cavpor3.0 | GCF_000151735.1 |
| Star-nosed mole (*Condylura cristata*) | ConCri1.0 | GCF_000260355.1 |
| Big brown bat (*Eptesicus fuscus*) | EptFus1.0 | GCF_000308155.1 |
| Horse (*Equus caballus*) | EquCab3.0 | GCF_002863925.1 |
| Western European hedgehog (*Erinaceus europaeus*) | EriEur2.0 | GCF_000296755.1 |
| Human (*Homo sapiens*) | GRCh38.p14 | GCF_000001405.40 |
| African savanna elephant (*Loxodonta africana*) | Loxafr3.0 | GCF_000001905.1 |
| Gray mouse lemur (*Microcebus murinus*) | Mmur_3.0 | GCF_000165445.2 |
| Prairie vole (*Microtus ochrogaster*) | MicOch1.0 | GCF_000317375.1 |
| House mouse (*Mus musculus*) | GRCm39 | GCF_000001635.27 |
| Etruscan shrew (*Suncus etruscus*) | mSunEtr1.pri.cur | GCF_024139225.1 |
| American pika (*Ochotona princeps*) | OchPri3.0 | GCF_000292845.1 |

| | | |
|---|---|---|
| Killer whale (*Orcinus orca*) | Oorc_1.1 | GCF_000331955.2 |
| Norway rat (*Rattus norvegicus*) | mRatBN7.2 | GCF_015227675.2 |
| Common shrew (*Sorex araneus*) | SorAra2.0 | GCF_000181275.1 |
| Smoky shrew (*Sorex fumeus*) | SorFum_2.1 | GCF_029834395.1 |
| Maritime shrew (*Sorex maritimensis*) | SorMar_1.0* | GCA_030324115.1* |
| Pig (*Sus scrofa*) | Sscrofa11.1 | GCF_000003025.6 |

***Table S2.*** *Conserved regions and accelerated region statistics based on the hg38 chromosomes (NCBI accession: GCA_000001405.15) for all shrew species, and the African elephant and big brown bat.*

| Chromosome | # 50 bp conserved regions | # ARs S. etruscus | # ARs S. cinereus | # ARs S. maritimensis | # ARs S. araneus | #ARs L. africana | #ARs E. fuscus |
|---|---|---|---|---|---|---|---|
| **1** | 53,160 | 1,494 | 548 | 2,183 | 259 | 67 | 2,870 |
| **2** | 62,618 | 2,638 | 315 | 1,262 | 145 | 362 | 1,942 |
| **3** | 45,064 | 1,131 | 365 | 1,317 | 139 | 177 | 1,253 |
| **4** | 28,793 | 962 | 156 | 615 | 104 | 73 | 1,266 |
| **5** | 38,505 | 983 | 219 | 1,393 | 212 | 38 | 1,475 |
| **6** | 32,985 | 1,358 | 325 | 871 | 144 | 93 | 422 |
| **7** | 31,500 | 988 | 124 | 939 | 124 | 158 | 915 |
| **8** | 26,899 | 986 | 233 | 512 | 67 | 86 | 1,166 |
| **9** | 29,383 | 778 | 154 | 735 | 73 | 178 | 436 |
| **10** | 26,248 | 2,080 | 153 | 841 | 126 | 77 | 928 |
| **11** | 30,470 | 1,056 | 244 | 1,667 | 260 | 65 | 1,144 |
| **12** | 27,793 | 1,237 | 344 | 1,371 | 197 | 28 | 175 |
| **13** | 18,667 | 536 | 121 | 395 | 39 | 190 | 324 |
| **14** | 24,584 | 805 | 80 | 806 | 73 | 10 | 343 |
| **15** | 20,174 | 1,143 | 104 | 816 | 111 | 54 | 1,146 |
| **16** | 21,988 | 651 | 187 | 1,138 | 117 | 70 | 1,890 |
| **17** | 26,095 | 978 | 166 | 1,600 | 181 | 15 | 228 |
| **18** | 16,292 | 355 | 77 | 348 | 30 | 54 | 1,685 |
| **19** | 13,331 | 457 | 142 | 1,575 | 126 | 41 | 768 |
| **20** | 13,448 | 421 | 120 | 609 | 70 | 35 | 51 |
| **21** | 4,008 | 103 | 19 | 120 | 14 | 24 | 467 |
| **22** | 6,175 | 214 | 76 | 468 | 32 | 9 | 45 |
| **TOTAL** | **598,180** | **21,354** | **4,272** | **21,581** | **2,643** | **1,904** | **20,939** |

***Table S3.*** *References for genes of interest in Table 4.*


Aigner, L., Arber, S., Kapfhammer, J. P., Laux, T., Schneider, C., Botteri, F., Brenner, H.-R., & Caroni, P. (1995). Overexpression of the neural growth-associated protein GAP-43 induces nerve sprouting in the adult nervous system of transgenic mice. Cell, 83(2), 269–278. https://doi.org/10.1016/0092-8674(95)90168-X

Chrysanthou, A., Ververis, A., & Christodoulou, K. (2022). ANO10 Function in Health and Disease. The Cerebellum, 22(3), 447–467. https://doi.org/10.1007/s12311-022-01395-3

Franchini, L., Stanic, J., Ponzoni, L., Mellone, M., Carrano, N., Musardo, S., Zianni, E., Olivero, G., Marcello, E., Pittaluga, A., Sala, M., Bellone, C., Racca, C., Di Luca, M., & Gardoni, F. (2019). Linking NMDA Receptor Synaptic Retention to Synaptic Plasticity and Cognition. IScience, 19, 927–939. https://doi.org/10.1016/j.isci.2019.08.036

Kimura, H., Fujita, Y., Kawabata, T., Ishizuka, K., Wang, C., Iwayama, Y., Okahisa, Y., Kushima, I., Morikawa, M., Uno, Y., Okada, T., Ikeda, M., Inada, T., Branko, A., Mori, D., Yoshikawa, T., Iwata, N., Nakamura, H., Yamashita, T., & Ozaki, N. (2017). A novel rare variant R292H in RTN4R affects growth cone formation and possibly contributes to schizophrenia susceptibility. Translational Psychiatry, 7(8), e1214–e1214. https://doi.org/10.1038/tp.2017.170

Latremoliere, A., Cheng, L., DeLisle, M., Wu, C., Chew, S., Hutchinson, E. B., Sheridan, A., Alexandre, C., Latremoliere, F., Sheu, S.-H., Golidy, S., Omura, T., Huebner, E. A., Fan, Y., Whitman, M. C., Nguyen, E., Hermawan, C., Pierpaoli, C., Tischfield, M. A., … Engle, E. C. (2018). Neuronal-Specific TUBB3 Is Not Required for Normal Neuronal Function but Is Essential for Timely Axon Regeneration. Cell Reports, 24(7), 1865-1879.e9. https://doi.org/10.1016/j.celrep.2018.07.029

Lee, Y. J., Jeong, Y. J., Kang, E. J., Kang, B. S., Lee, S. H., Kim, Y. J., Kang, S. S., Suh, S. W., & Ahn, E. H. (2023). GAP-43 closely interacts with BDNF in hippocampal neurons and is associated with Alzheimer's disease progression. Frontiers in Molecular Neuroscience, 16. https://doi.org/10.3389/fnmol.2023.1150399

Li, R., Meng, M., Chen, Y., Pan, T., Li, Y., Deng, Y., Zhang, R., Tian, R., Xu, W., Zheng, X., Gong, F., Liu, J., Tang, H., Ding, X., Tang, Y., Annane, D., Chen, E., Qu, H., & Li, L. (2023). ATP-citrate lyase controls endothelial gluco-lipogenic metabolism and vascular inflammation in sepsis-associated organ injury. Cell Death & Disease, 14(7), 401. https://doi.org/10.1038/s41419-023-05932-8

Long, H., Zhu, X., Yang, P., Gao, Q., Chen, Y., & Ma, L. (2013). Myo9b and RICS Modulate Dendritic Morphology of Cortical Neurons. Cerebral Cortex, 23(1), 71–79. https://doi.org/10.1093/cercor/bhr378

Ohkubo, Y., Uchida, A. O., Shin, D., Partanen, J., & Vaccarino, F. M. (2004). Fibroblast Growth Factor Receptor 1 Is Required for the Proliferation of Hippocampal Progenitor Cells and for Hippocampal Growth in Mouse. The Journal of Neuroscience, 24(27), 6057–6069. https://doi.org/10.1523/JNEUROSCI.1140-04.2004

Okur, V., Cho, M. T., Henderson, L., Retterer, K., Schneider, M., Sattler, S., Niyazov, D., Azage, M., Smith, S., Picker, J., Lincoln, S., Tarnopolsky, M., Brady, L., Bjornsson, H. T., Applegate, C., Dameron, A., Willaert, R., Baskin, B., Juusola, J., & Chung, W. K. (2016). De novo mutations in CSNK2A1 are associated with neurodevelopmental abnormalities and dysmorphic features. Human Genetics, 135(7), 699–705. https://doi.org/10.1007/s00439-016-1661-y

Palma, F. Di, Holme, R. H., Bryda, E. C., Belyantseva, I. A., Pellegrino, R., Kachar, B., Steel, K. P., & Noben-Trauth, K. (2001). Mutations in Cdh23, encoding a new type of cadherin, cause stereocilia disorganization in waltzer, the mouse model for Usher syndrome type 1D. Nature Genetics, 27(1), 103–107. https://doi.org/10.1038/83660

Reinhard, J. R., Kriz, A., Galic, M., Angliker, N., Rajalu, M., Vogt, K. E., & Ruegg, M. A. (2016). The calcium sensor Copine-6 regulates spine structural plasticity and learning and memory. Nature Communications, 7(1), 11613. https://doi.org/10.1038/ncomms11613

Roux, I., Safieddine, S., Nouvian, R., Grati, M., Simmler, M.-C., Bahloul, A., Perfettini, I., Le Gall, M., Rostaing, P., Hamard, G., Triller, A., Avan, P., Moser, T., & Petit, C. (2006). Otoferlin, Defective in a Human Deafness Form, Is Essential for Exocytosis at the Auditory Ribbon Synapse. Cell, 127(2), 277–289. https://doi.org/10.1016/j.cell.2006.08.040

Roychaudhuri, R., Atashi, H., & Snyder, S. H. (2023). Serine Racemase mediates subventricular zone neurogenesis via fatty acid metabolism. Stem Cell Reports, 18(7), 1482–1499. https://doi.org/10.1016/j.stemcr.2023.05.015

Seidler, N. W. (2013). GAPDH and Intermediary Metabolism (pp. 37–59). https://doi.org/10.1007/978-94-007-4716-6_2

Wong, J. M., Folorunso, O. O., Barragan, E. V., Berciu, C., Harvey, T. L., Coyle, J. T., Balu, D. T., & Gray, J. A. (2020). Postsynaptic Serine Racemase Regulates NMDA Receptor Function. The Journal of Neuroscience, 40(50), 9564–9575. https://doi.org/10.1523/JNEUROSCI.1525-20.2020

Yamauchi, T., Nio, Y., Maki, T., Kobayashi, M., Takazawa, T., Iwabu, M., Okada-Iwabu, M., Kawamoto, S., Kubota, N., Kubota, T., Ito, Y., Kamon, J., Tsuchida, A., Kumagai, K., Kozono, H., Hada, Y., Ogata, H., Tokuyama, K., Tsunoda, M., … Kadowaki, T. (2007). Targeted disruption of AdipoR1 and AdipoR2 causes abrogation of adiponectin binding and metabolic actions. Nature Medicine, 13(3), 332–339. https://doi.org/10.1038/nm1557

York, N. S., Sanchez-Arias, J. C., McAdam, A. C. H., Rivera, J. E., Arbour, L. T., & Swayne, L. A. (2022). Mechanisms underlying the role of ankyrin-B in cardiac and neurological health and disease. Frontiers in Cardiovascular Medicine, 9. https://doi.org/10.3389/fcvm.2022.964675

Zhao, J., Chen, J., Li, M., Chen, M., & Sun, C. (2020). Multifaceted Functions of CH25H and 25HC to Modulate the Lipid Metabolism, Immune Responses, and Broadly Antiviral Activities. Viruses, 12(7), 727. https://doi.org/10.3390/v12070727
